# On Weighted K-Mer Dictionaries

**DOI:** 10.1101/2022.05.23.493024

**Authors:** Giulio Ermanno Pibiri

## Abstract

We consider the problem of representing a set of *k*-mers and their abundance counts, or weights, in compressed space so that assessing membership and retrieving the weight of a *k*-mer is efficient. The representation is called a *weighted dictionary* of *k*-mers and finds application in numerous tasks in Bioinformatics that usually count *k*-mers as a pre-processing step. In fact, *k*-mer counting tools produce very large outputs that may result in a severe bottleneck for subsequent processing.

In this work we extend the recently introduced SSHash dictionary (Pibiri, *Bioinformatics* 2022) to also store compactly the weights of the *k*-mers. From a technical perspective, we exploit the order of the *k*-mers represented in SSHash to encode *runs* of weights, hence allowing (several times) better compression than the empirical entropy of the weights. We also study the problem of reducing the number of runs in the weights to improve compression even further and illustrate a lower bound for this problem. We propose an efficient, greedy, algorithm to reduce the number of runs and show empirically that it performs well, i.e., very similarly to the lower bound. Lastly, we corroborate our findings with experiments on real-world datasets and comparison with competitive alternatives. Up to date, SSHash is the only *k*-mer dictionary that is exact, weighted, associative, fast, and small.

## 1 Introduction

Recent advancements in the so-called Next Generation Sequencing (NGS) technology made possible the availability of very large collections of DNA. However, before being able to actually analyze the data at this scale, efficient methods are required to index and search such collections. One popular strategy to address this challenge is to consider short substrings of fixed length *k*, known as *k*-mers. Software tools based on *k*-mers are predominant in Bioinformatics and they have found applications in genome assembly [3, 13], variant calling [15, 37], pan-genome analysis [2, 18], meta-genomics [38], sequence comparison [31, 33, 34], just to name a few but noticeable ones.

For several such applications it is important to quantify how many times a given *k*-mer is present in a DNA database. In fact, many efficient *k*-mer counting tools have been developed for this task [8, 21, 16, 30]. The output of these tools is a table where each distinct *k*-mer in the database is associated to its abundance count, or *weight*. The weights are either exact or approximate. (In this work, we focus on exact weights.) These genomic tables are usually very large and take several GBs – in the range of 40-80 bits/*k*-mer or more according to recent experiments [12, 19, 21]. Therefore, the tables should be compressed effectively while permitting efficient random access queries in order to be useful for on-line processing tasks. This is precisely the goal of this work. We better formalize the problem as follows.

Let 𝒦 be the set of the *n* distinct *k*-mers extracted from a given DNA string. In particular, 𝒦 can be regarded as a set of *n* pairs ⟨*g, w*(*g*) ⟩, where *g* is a *k*-mer and *w*(*g*) is the *weight* of *g*. Our objective is to build a compressed, weighted, dictionary for 𝒦, i.e., a data structure representing the *k*-mers *and* weights of 𝒦 in compressed space such that it is efficient to check the *exact* membership of *g* to 𝒦 and, if *g* actually belongs to 𝒦, retrieve *w*(*g*).

In our previous investigation on the problem, we proposed a *sparse and skew hashing* scheme for *k*-mers (SSHash, henceforth) [22] – a compressed dictionary that relies on *k*-mer *minimizers* [31] and *minimal perfect hashing* [23, 24] to support fast membership (in both random and streaming query modality) in succinct space. However, we did not consider the weights of the *k*-mers. In this work, therefore, we enrich the SSHash data structure with the weight information. The main practical result is that, by exploiting the *order* of the *k*-mers represented in SSHash, the compressed exact weights take only a small extra space on top of the space of SSHash. This extra space is proportional to the number of *runs* (maximal sub-sequences formed by all equal symbols) in the weights and not proportional to the number of distinct *k*-mers. As a consequence, the weights are represented in a much smaller space than the empirical entropy lower bound.

We also study the problem of reducing the number of runs in the weights and model it as a *graph covering* problem, for which we give an expected linear-time algorithm. The optimization algorithm effectively reduces the number of runs, hence improving space even further, and almost matches the best possible reduction according to a lower bound.

When empirically compared to other weighted dictionaries that can be either somewhat smaller but much slower or much larger, SSHash embodies a robust trade-off between index space and query efficiency.

## 2 Related Work

A solution to the weighted *k*-mer dictionary problem can be obtained using the popular FM-index [11]. The FM-index represents the original DNA string taking the Burrows-Wheeler transform (BWT) [5] of the string. Reporting the weight of a *k*-mer is solved using the *count* operation of the FM-index which involves *O*(*k*) rank queries over the BWT.

Another solution using the BWT is the so-called BOSS data structure [4] that is a succinct representation of the *de Bruijn* graph of the input – a graph where the nodes are the *k*-mers and the edges model the overlaps between the *k*-mers. The BOSS data structure has been recently enriched with the weights of the *k*-mers [12], by delta-encoding the weights on a spanning branching of the graph. Since consecutive *k*-mers often have equal (or very similar) weights, good space effectiveness is achieved by this technique.

Other solutions, instead, rely on hashing for faster query evaluation compared to BWT-based indexes. For example, both deBGR [19] and Squeakr [21] uses a *counting quotient filter* [20] to store the *k*-mers and the weights. They can either return approximate weights, i.e., wrong answers with a prescribed (low) probability, for better space usage of exact weights at the price of more index space. In any case, the memory consumption of these solutions is not competitive with that of BWT-based ones as they do not employ sophisticated compression techniques and were designed for other purposes, e.g., dynamic updates.

A closely related problem is that of realizing *maps* from *k*-mers to weights, i.e., data structures that do *not* explicitly represent the *k*-mers and so return arbitrary answers for out-of-set keys. In the context of this work, we distinguish between such approaches, *maps*, and dictionaries that instead represent *both* the *k*-mers and the weights. Besides minimal perfect hashing [23, 24], some efficient maps have been proposed and tailored specifically for genomic counts, such as based on *set-min sketches* [36] and *compressed static functions* (CSFs) [35]. These proposals leverage on the repetitiveness of the weights (low-entropy distributions) to obtain very compact space.

Lastly in this section, we report that other works [17, 14] considered the multi-document version of the problem studied here, that is, how to retrieve a vector of weights for a query *k*-mer, where each component of the vector represents the weight of the *k*-mer in a distinct document. Also such count vectors are usually very “regular” (or can be made so by introducing some approximation) [17] and present runs of equal symbols that can be compressed effectively with *run-length encoding* (RLE).

## 3 Representing Runs of Weights

In this section we describe the compression scheme for the weights that we use in SSHash. Recall that we indicate with 𝒦 the set of *n* distinct ⟨*k*-mer, weight⟩ = ⟨*g, w*(*g*) ⟩ pairs, that we want to store in a dictionary. We first highlight the main properties of SSHash that we are going to exploit in the following to obtain good space effectiveness for the weights. (For all the other details concerning the SSHash index, we point the interested reader to our previous work [22].)

From a high-level perspective, SSHash implements the function *h* : Σ^*k*^ → {0, 1, …, *n*}, where *n* =|𝒦| and Σ^*k*^ is the whole set of *k*-length strings over the DNA alphabet Σ = {`A`, `C`, `G`, `T`}. In particular, *h*(*g*) is a unique value 1 ≤ *i* ≤ *n* if *g* ∈ 𝒦; or *h*(*g*) = 0 otherwise, i.e., if *g* ∉ 𝒦. In other words, SSHash serves the same purpose of a minimal and perfect hash function (MPHF) [24] for 𝒦 but, unlike a traditional MPHF, SSHash *rejects* alien *k*-mers. This is possible because the *k*-mers of 𝒦 are actually represented in SSHah whereas the space of a traditional MPHF does not depend on the input keys.

The value *i* = *h*(*g*) for the *k*-mer *g* ∈ 𝒦 is the handle of *g*, or its “hash” code. The hash codes can be used to associate some satellite information to the *k*-mers such as, for example, the weights themselves using an array *W* [1..*n*] where *W* [*h*(*g*)] = *w*(*g*). The crucial property of SSHash in which we are interested is that the function *h preserves the relative order* of the *k*-mers, that is: if *g*_1_[1..*k*] and *g*_2_[1..*k*] are two *k*-mers with *g*_1_[2..*k*] = *g*_2_[1..*k* −1] (i.e., *g*_2_ comes immediately after *g*_1_ in a string), then *h*(*g*_2_) = *h*(*g*_1_) + 1. Therefore, consecutive *k*-mers, i.e., those sharing an overlap of *k* − 1 symbols, are also given consecutive hash codes. This is achieved in SSHash by pre-processing the input set 𝒦 into a so-called *spectrum-preserving string set* (or SPSS) 𝒮 – a collection of strings 𝒮 = {𝒮_1_, …, 𝒮_*m*_} where each *k*-mer of 𝒦 appears exactly once. We omit the details here on how the collection 𝒮 can be built; we only report that there are efficient algorithms for this purpose that also try to minimize the total number of symbols in 𝒮, i.e., the quantity 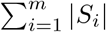. One such algorithm is the UST algorithm [29] that we also use to prepare the input for SSHash.

Therefore, once an order 𝒮_1_, …, 𝒮_*m*_ for the strings of 𝒮 is fixed, then also an order *i* = 1, …, *n* for the *k*-mers *g*_*i*_ is uniquely determined. Let *W* [1..*n*] be the sequence of weights in this order. Then, we have: *h*(*g*_*i*_) = *i* and *W* [*i*] = *w*(*g*_*i*_), for *i* = 1, …, *n*.

This order-preserving behavior of *h* induces a property on the sequence of weights *W* [1..*n*] that significantly aids compression: *W* contains *runs*, i.e., maximal sub-sequences of *equal weights*. This is so because consecutive *k*-mers are very likely to have the same weight due to the high specificity of the strings. This a known fact, also observed in prior work [12, 17, 35]. Here, we are exploiting the order of the *k*-mers given by SSHash to preserve the natural order of the weights in *W*. Note that this cannot be achieved by approximate schemes that do not represent the *k*-mers themselves, like a generic MPHF or a CSF. Even if the *k*-mers were available, those schemes are unable to assign consecutive hashes to consecutive *k*-mers, actually shuffling the weights at random and, thus, making *W* very difficult to compress.

It is standard to represent a sequence *W* featuring *r* runs of equal symbols using *run-length encoding* (RLE), i.e., *W* is modeled as a sequence of run-length pairs *RLW* = ⟨*w*_1_, 𝓁_1_⟩ ⟨*w*_2_, 𝓁_2_⟩ … ⟨*w*_*r*_, 𝓁_*r*_⟩ where *w*_*i*_ and 𝓁_*i*_ are, respectively, the value of the run and the length of the *i*-th run in *W*. Figure 1 shows an example of *RLW* for a collection 𝒮 with 4 weighted strings.

**Figure 1.**
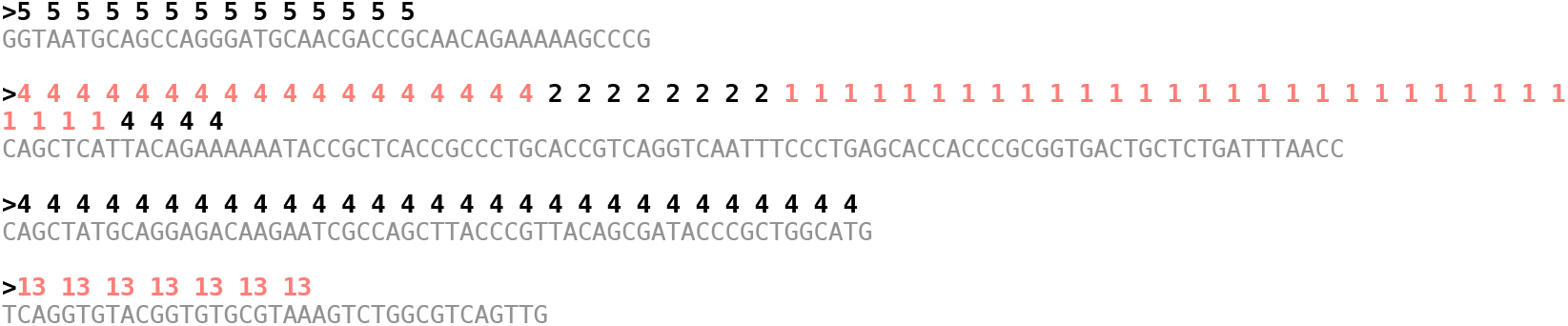
An example collection 𝒮 of 4 weighted sequences (for *k* = 31) drawn from the genome of *E. Coli* (Sakai strain). With alternating colors we render the change of weight in the runs. There are 111 *k*-mers in the example but just 6 runs in the weights: *RLW* = ⟨5, 14⟩ ⟨4, 18⟩ ⟨2, 8⟩ ⟨1,31⟩ ⟨4, 33⟩ ⟨13, 7⟩. Note that a run can cross the boundary between two (or more) sequences, as it happens for the run ⟨4, 18⟩ which covers completely the third but also the part of the second sequence.

### Encoding RLW

Let *D* be the set of all distinct *w*_*i*_ in *RLW*. Clearly, *r* ≥ |𝒟| as we must have at least one run per distinct weight. We store 𝒟 using |𝒟|⌈log_2_ *max*⌉ bits where *max* ≥ 1 is the largest *w*_*i*_. We use 𝒟 to uniquely represent each *w*_*i*_ in *RLW* with ⌈log_2_|𝒟|⌉ bits. Since runs are maximal sub-sequences in *W* by definition, then *w*_*i*_ ≠ *w*_*i*+1_ for every *i* = 1, …, *r* − 1 (adjacent weights must be different). Then we take the prefix-sums of the sequence 0, 𝓁_1_, …, 𝓁_*r*−1_ into an array *L*[1..*r*] and encode it with Elias-Fano [9, 10]^1^. By construction we have that 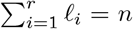 since the runs must cover the whole set of *k*-mers. So the largest element in *L* is actually *n* − 𝓁_*r*_ and we spend at most *r*⌈log_2_(*n/r*)⌉ + 2*r* + *o*(*r*) bits for *L*. Summing up, we spend at most

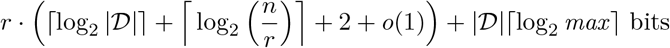

for representing *RLW* on top of the space of SSHash. In conclusion, the weights are represented in space proportional to the number of runs in *W* (i.e., *r* = |*RLW*|) and *not* proportional to the number of *k*-mers, which is *n*. As a consequence, this space is likely to be considerably less than the empirical entropy *H*_0_(*W*) as we are going to see with the experiments in Section 5.

To retrieve the weight *w*(*g*) from *i* = *h*(*g*), all that is required is to identify the run containing *i*. This operation is done in *O*(log(*n/r*)) time with a predecessor query over *L* given that we represent *L* with Elias-Fano. If the identified run is the *j*-th run in *W*, then *w*_*j*_ is retrieved in *O*(1) from 𝒟.

## 4 Reducing the Number of Runs

In Section 3 we presented an encoding scheme for the *k*-mer weights whose space is proportional to the *number of runs* in the sequence of weights *W*. Therefore, in this section we consider the problem of reducing the number of runs in the weights to optimize the space of the encoding.

### Rules of the Game

We assume that the strings in 𝒮 are *atomic* entities: it is not allowed to partition them into sub-strings (e.g., in correspondance of the runs of weights in the strings). In fact, since the strings are obtained by the UST algorithm [29] with the purpose of *minimizing* the number of nucleotides as we explained in Section 3, breaking them will lead to an increased space usage for the *k*-mers, actually dwarfing any space-saving effort spent for the weights. With this constraint specified, there are only *two* degrees of freedom that can be exploited to obtain better compression for *W* : (1) the *order* of the strings, and (2) the *orientation* of the strings. Altering 𝒮 using these two degrees of freedom does not affect the correctness nor the (relative) order-preserving property of the function *h* : Σ^*k*^ → {0, 1, …, *n*} implemented by SSHash. In fact, as evident from our description in Section 3, the output of *h* will still be {1, …, *n*} as the *k*-mers themselves do *not* change (even when taking reverse-complements into account as they are considered to be identical). What changes is just the absolute order of the *k*-mers as a consequence of permuting the order of the strings {*S*_1_, …, *S*_*m*_} in 𝒮.

Therefore, our goal is to permute the order of the strings in 𝒮 and possibly change their orientations to reduce the number of runs in *W*. We now consider an illustrative example to motivate why both these two operations – those of changing the order and orientation of a string – are important to reduce the number of runs. Refer to Figure 2a which shows an example collection of *m* = |𝒮| = 4 weighted strings (for *k* = 3). Applying the permutation *π* = [1, 4, 2, 3] as shown in Figure 2b reduces the number of runs by 1 because the run at the junction of string 4 and 2 can be glued. Lastly, applying the *signed* permutation *π* = [+1, +4, −2, +3] as in Figure 2c reduces the number of runs by 3, which is the best possible. Our objective is to compute such a signed permutation *π* for an input collection of strings, in order to permute 𝒮 as shown in Algorithm 1.

**Figure 2.**
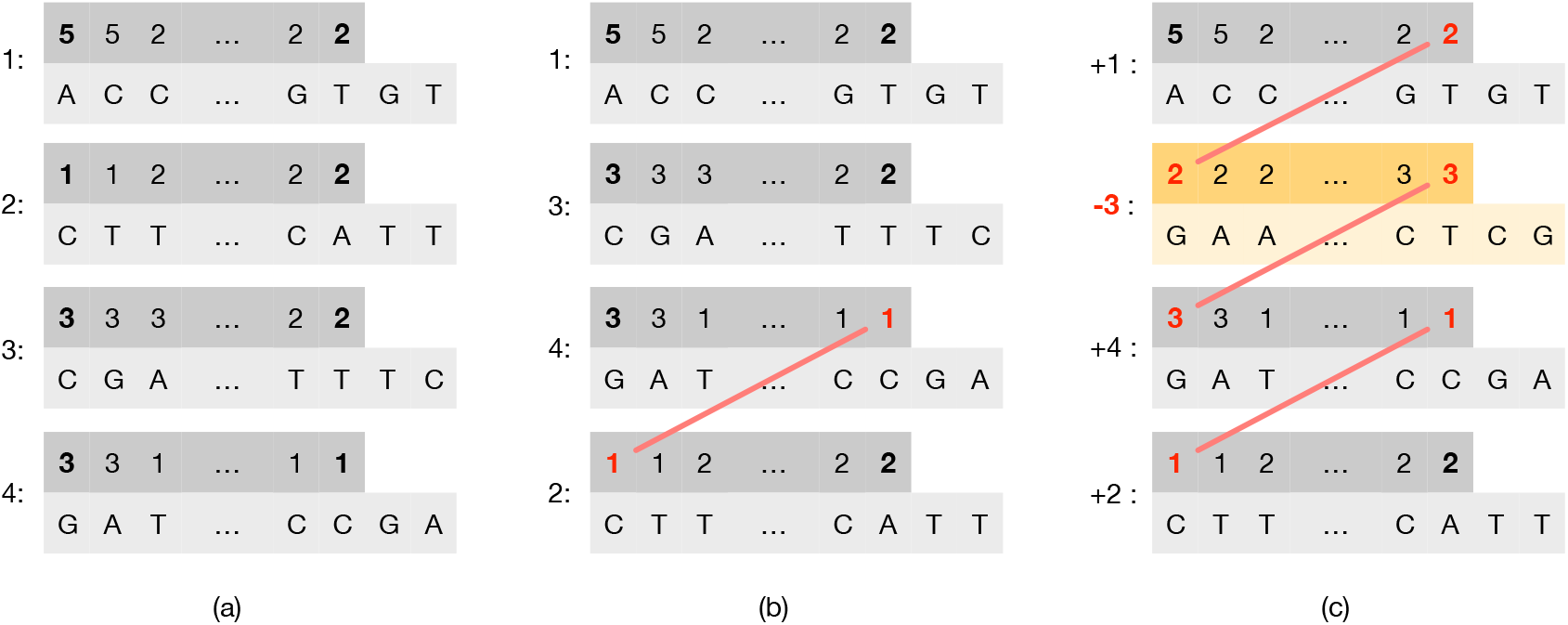
In (a), an example input collection 𝒮 of *m* =|𝒮| = 4 weighted strings (for *k* = 3), where the end-point weights are highlighted in bold font. In (b), the order of the strings is changed according to the permutation *π* = [1, 4, 2, 3] and, as a result, the number of runs is reduced by 1 (the last run in string 4 is glued with the first run of string 2). Lastly, in (c), it is shown that changing the orientation of string 3 (taking the reverse complement of the string and reversing the order of the *k*-mer weights) makes it possible to glue other two runs. Given that reducing the number of runs by *m* − 1 is the best achievable reduction, the number of runs in (c) is therefore the minimum for the original collection in (a).

#### Algorithm 1

The algorithm permute takes as input a collection 𝒮 = {*S*_1_, …, *S*_*m*_} of weighted strings and a signed permutation *π* and returns the permuted collection 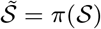. The function reverse takes the reverse-complement of a string and reverse its weights.

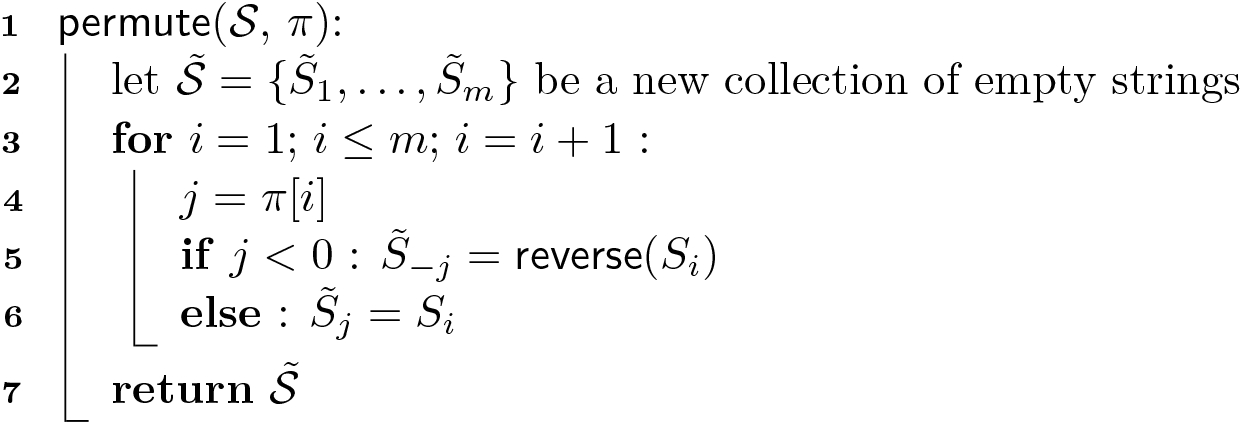

Figure 2 also suggests that the final result *π* solely depends on the weight of the first and last *k*-mer of each sequence – which we call the *end-point* weights (or just end-points) of a sequence – and not on the other weights nor the nucleotide sequences. Therefore, it is useful to model an input collection 𝒮 using a graph, defined as follows.

#### Definition 1

`(End-point Weight Graph)`. Given a collection of weighted sequences 𝒮, let *G* be a graph where:

▬ There is a node *u* for each sequence of 𝒮 and *u* has two sides – a *left* and a *right* side – respectively labelled with the end-point weights of the sequence.
▬ There is an edge between any two distinct nodes *u* and *v* that have a side with the same weight.

The graph *G* is called the *end-point weight graph* for 𝒮 and indicated with *ewG*(𝒮).

In the following, we indicate a node *u* in *ewG*(𝒮) using the identifier (*id*) of the corresponding sequence of 𝒮. Also, we associate to *u* a *sign* ∈ {−1, +1} (or orientation), indicating whether the sequence should be reverse-complemented. In summary, a node *u* in *ewG*(𝒮) is the 4-tuple (*id, left, right, sign*).

#### Definition 2

`(Oriented Path)`. An *oriented* path in *ewG*(𝒮) is either a single node (singleton path) or a sequence of nodes (*u*_1_ → … → *u*_𝓁_) where each consecutive pair of nodes *u*_*i*_ → *u*_*i*+1_ is oriented in such a way that *u*_*i*_.*right* = *u*_*i*+1_.*left*, for any 1 ≤ *i <* 𝓁.

Since we will be interested only in oriented paths, we just refer to them as “paths”. For ease of notation, we will indicate a path in our examples as a sequence of signed numbers (*i*_1_→ … →*i*_𝓁_) where each number represents a node’s *id* and its sign represents the node’s *sign*. The first and the last node in the path are called, respectively, the *front* and the *back* of the path. The weights *front*.*left* and *back*.*right* are the two end-points of the path.

Given this graph model, it follows that the problem of finding a signed permutation *π* for 𝒮 is equivalent to that of computing a (disjoint-node) *path cover C* for *ewG*(𝒮), i.e., a set of paths in *ewG*(𝒮) that visit *all* the nodes and where each node belongs to *exactly one* path. In fact note that, given a cover *C* for *ewG*(𝒮), there is a linear-time reduction from *C* to *π* as illustrated in Algorithm 2. Since the cover *C* is a disjoint-node path cover, the correctness of the algorithm is immediate as well as its complexity of Θ(*m*).

Figure 3 illustrates the same example of Figure 2 but with end-point weight graphs. In Figure 3b we would obtain a cover *C* = {(+1 → −3 → +4 → +2)} formed by a single path. In this case the permutation *π*, following Algorithm 2, would be *π*[1] = +1, *π*[3] = −2, *π*[4] = +3, and *π*[2] = +4. This is indeed the same permutation discussed in Figure 2c. Another example: for the graph in Figure 3c, the cover would be *C* = {(+2 → −3 → +4), (+1)} and the permutation *π* would be *π*[2] = +1, *π*[3] = −2, *π*[4] = +3, and *π*[1] = +4.

**Figure 3.**
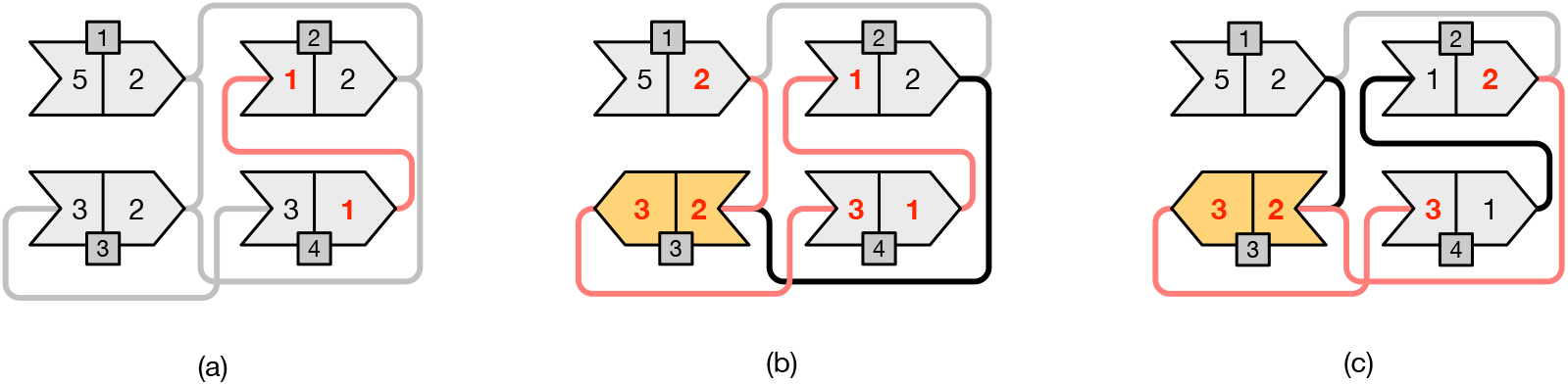
The same example of Figure 2 but modeled using end-point weight graphs. Each node is represented using an arrow-like shape with two-matching sides. Only opposite sides having the same weight can be matched. The numbers inside the shapes represent the end-point weights; the extra darker square contains the node identifier. An arrow oriented from *left-to-right* models a node with *positive* sign; vice versa, an arrow oriented from *right-to-left* models a node with *negative* sign. Gray edges represent edges that *cannot* be traversed without changing the orientation of one of the two connected nodes. Black edges represent edges that can be traversed. Lastly, we highlight in red the edges that belong to paths in a graph cover. The example in (a) corresponds to that of Figure 2b where no node has changed orientation and, therefore, we have three paths in the cover: (+4 → +2), (+3), and (+1). Other two different covers are shown in (b) and (c). In (b) the cover contains the single path (+1 → −3 → +4 → +2) and corresponds to the example of Figure 2c where the node 3 was changed orientation from + to − (shown in yellow color). In (c) the cover contains the two paths (+2 → −3 → +4) and the singleton path (+1).

#### Algorithm 2

The algorithm reduce takes as input a path cover *C* computed for *ewG*(𝒮) and returns the corresponding signed permutation *π*. The complexity of the algorithm is Θ(*m*) since the number of nodes in *ewG*(𝒮) is *m* and each node appears exactly once in *C*.

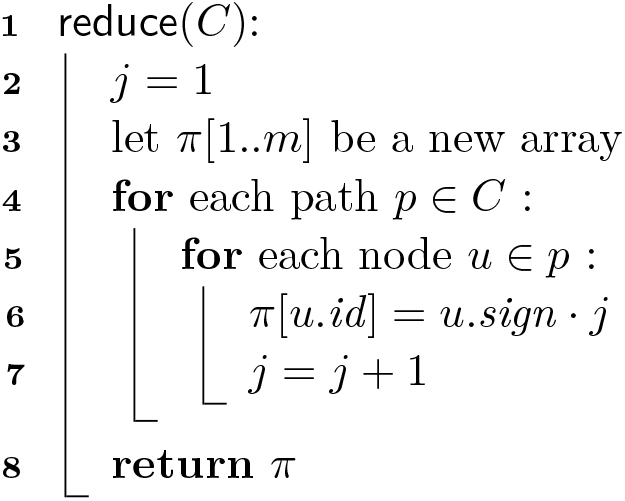

#### 4.1 A Lower Bound to the Number of Runs

We showed that changing the order and orientation of the strings in 𝒮 can reduce the number of runs in the weights. The crucial question is: by how much? We are interested in deriving a lower bound to the number of runs achievable after applying the signed permutation *π* to 𝒮. Since we modeled the problem of computing *π* as the problem of finding a path cover *C* for *ewG*(𝒮), we reason in terms of *ewG*(𝒮) and *C*.

Let |*C*| be the number of paths in the cover *C*. Let *r*_*i*_ be the number of runs in *S*_*i*_ and let *R* be the total number of runs, i.e., 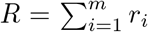. Then there are at least *R* – *m* runs in 𝒮 *regardless* the order of the sequences. Therefore, a straightforward lower bound to the number of runs would be max {|𝒟 |, *R* − *m* + 1}. This lower bound assumes (very optimistically) that we are able to obtain a cover with a single path, i.e., |*C*| = 1, hence reducing the total number of runs *R* by *m* − 1 which is the best reduction achievable with *m* sequences. Note, however, that the bound cannot be lower than |𝒟| – the number of distinct weights in the input – because, clearly, there must be at least one run per distinct weight value. We would like to improve the bound max {|𝒟|, *R* − *m* + 1} knowing that, in general, we could not be able to form one single path.

We observe that the final number of runs *r* in the permuted 𝒮 will be equal to *R* − *m* + |*C*|. In fact, every path in *C* must begin (resp. end) with a node whose left side (resp. right side) cannot be glued with any other path’s side. Therefore, a new run begins with the first node of every path. Since we wish to minimize the quantity *R* − *m* + |*C*|, and considering that *R* − *m* is constant for a given 𝒮, it follows that the problem reduces to that of minimizing |*C*|, the number of paths in the cover. In other words, the problem of minimizing the number of runs *r* is equivalent to that of finding a minimum-cardinality path cover *C* for *ewG*(𝒮).

Therefore our strategy is to give a lower bound to the number of paths |*C*|. To do so, we compute the number of end-point weights, say *n*_*e*_, that must appear as end-points of the paths (left side of the front node of a path, or right side of the back node of a path). Since a path has exactly two end-points, then it follows that |*C*| ≥ ⌈*n*_*e*_*/*2⌉ and, in turn, that the final number of runs *r* in *π*(𝒮) is *at least R* − *m* + ⌈*n*_*e*_*/*2⌉ ≥ max{|𝒟|, *R* − *m* + 1}.

Since we now focus on the end-point weights of the nodes, in this section we will denote a node (*id, left, right, sign*) just by its weights (*left, right*). We start with a preliminary Lemma and a Definition. (See Appendix A for all the proofs omitted from the section.)

##### Lemma 3.

Let us consider *d >* 0 equal nodes (*w, x*). If *d* is even, then the *d* nodes originate a path of either end-points (*w, w*) or (*x, x*). If *d* is odd, then the path has end-points (*w, x*).

##### Definition 4

`(Incidence Set)`. Given the weight *w*, a set *I*_*w*_ of nodes where *w* appears as end-point is called an *incidence set* for *w*. Let *n*(*I*_*w*_) be the number of times *w* appears in the nodes of *I*_*w*_. Note that *n*(*I*_*w*_) ≥ |*I*_*w*_| because there could be nodes (*w, w*) in *I*_*w*_.

##### Example 5.

The sets

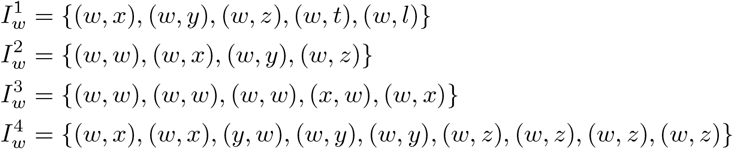

are four different incidence sets for *w* with 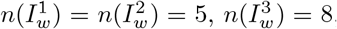, and 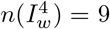. The set {(*w, x*), (*x, y*)}, instead, is not an incidence set for *w* because the node (*x, y*) does not have *w* as an end-point.

Next, we give the following central Lemma that will help in counting the number of weights that must appear as end-points of the paths in *C*.

##### Lemma 6.

Given an incidence set *I*_*w*_, if *n*(*I*_*w*_) is *odd* then only one path will contain *w* as end-point among all the paths that can be created from the nodes in *I*_*w*_.

##### Example 7.

Let us consider an example for Lemma 6. The sets 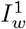 and 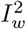 in Example 5 are both canonical since all other weights different from *w* are distinct, except for one single node (*w, w*) in 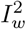. It is then easy to see that no matter what paths are created, there will always be one extra node that will remain alone. The set 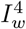 in Example 5, with 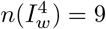, is not canonical instead and we have: *D*_*x*_ = {(*w, x*), (*w, x*)}, *D*_*y*_ = {(*y, w*), (*w, y*), (*w, y*)}, and *D*_*z*_ = {(*w, z*), (*w, z*), (*w, z*), (*w, z*)}, with *d*_*x*_ =|*D*_*x*_| = 2, *d*_*y*_ = |*D*_*y*_| = 3, and *d*_*z*_ = |*D*_*z*_| = 4. Since *d*_*x*_ = 2, then the nodes in *D*_*x*_ can be collapsed into either (*w, w*) or (*x, x*). Since *d*_*y*_ = 3, then the nodes in *D*_*y*_ can be collapsed to (*w, y*). Lastly, since *d*_*z*_ = 4, the nodes in *D*_*z*_ can be collapsed to either (*w, w*) or (*z, z*). Again, regardless the choices done, 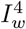 can always be reduced to a canonical set.

Let now *eW* be the set of the distinct end-point weights of 𝒮. For every weight *w* ∈ *eW*, let 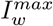 be the incidence set of *maximum-cardinality*, i.e., the one obtained by considering *all* the nodes in *ewG*(𝒮). We partition *eW* into three disjoint sets, *eW*_*odd*_, *eW*_*even*_, and *eW*_*equal*_. Based on the properties of 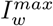, *w* belongs to one of the three sets as follows.

▬ If 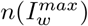 is odd, then *w* ∈ *eW*_*odd*_. In this case, we are sure by Lemma 6 that *w* will appear as end-point of some path in the cover.
▬ If all the nodes in 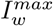 are equal to (*w, w*), then *w* ∈ *eW*_*equal*_. In this case 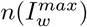 will always be even and, for Lemma 6, *w* will appear *twice* as end-points of the same path in the cover.
▬ If 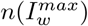 is even but the nodes in 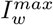 are *not* all equal to (*w, w*), then *w* ∈ *eW*_*even*_. In this case, we cannot be sure that *w* will *not* appear as end-point. If 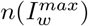 is even *and* 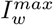 contains *all distinct* nodes, then *w* will not appear. But if 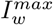 contains two identical nodes, say of the form (*w, x*), then *w* may or may not appear. In fact, as already noted, depending on whether a path is created as (*w, x*) → (*x, w*) or (*x, w*) → (*w, x*), *w* will appear (in the first case) or not (in the second case). The set 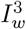 from Example 5 illustrates this case.

##### Lemma 8.

|*W*_*odd*_| is even.

Therefore, we obtain the following Corollary.

##### Corollary 9.

The number of paths in *C* is at least (2|*eW*_*equal*_| + |*eW*_*odd*_|)*/*2.

**Proof**. Immediate by first applying Lemma 6 to the maximal incidence sets, then noting that each each path has exactly two end-points, and lastly taking into account the cardinality of *eW*_*odd*_ for Lemma 8. ◄

Let us now compute the lower bound of Corollary 9 on some example graphs.

##### Example 10.

First, we re-consider the example graph from Figure 2. In that example we have a set of end-point weights *eW* = {1, 2, 3, 5}, where 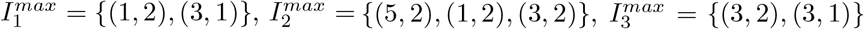, and 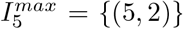. Since 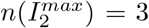 and 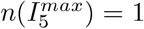, then *eW*_*odd*_ = {2, 5} and *eW*_*equal*_ = ∅. Therefore we sure that both weights 2 and 5 will appear as end-point weights of some paths in the cover. For Corollary 9, any path cover will contain at least (2|*eW*_*equal*_| + |*eW*_*odd*_|)*/*2 = (2 · 0 + 2)*/*2 = 1 path. Indeed the cover shown in Figure 2b contains one single path and so, it is optimal, whereas the cover in Figure 2c contains 2 paths and is not optimal.

##### Example 11.

We now consider the larger example in Figure 4 for a graph with 16 nodes. In this example we have a set of end-point weights *eW* = {1, 2, 3, 4, 7, 8, 13} and the incidence sets are as follows: 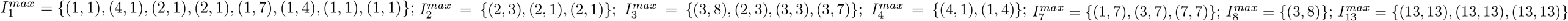. Since we have 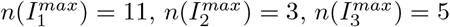, and 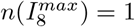, then *eW*_*odd*_ = {1, 2, 3, 8} and |*eW*_*odd*_| = 4 which is even (Lemma 8). Therefore we are sure that the weights 1, 2, 3, and 8 will appear as end-points of some paths in the cover. The incidence set 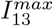 is made of nodes (13, 13), so *eW*_*equal*_ = {13} and |*eW*_*equal*_| = 1. Also in this case we are sure the weight 13 will appear (twice) as end-point of some path. (*eW*_*even*_ = {4, 7}.) For Corollary 9 we derive that a path cover for the example graph will contain at least (2 · 1 + 4)*/*2 = 3 paths. Figure 4 shows two different path covers for the same graph, that are *C*_*a*_ = {(+8 → +3 → +7 → +13 → −12 → +11 → −4 → +6 → +10 → +9 → +1 → −5), (+2), (+14 → +15 → +16)} and *C*_*b*_ = {(+1 → −3 → −8 → +9 → +10 → −5 → +6 → +7 → +12 → +2), (+4 → +11), (+13), (+14 → +15 → +16)}. The cover *C*_*a*_ is optimal for the lower bound as is contains 3 paths; the cover *C*_*b*_ contains 4 paths (1 more than necessary). It is not difficult to see that we cannot find a path cover with less than 3 paths for the graph in Figure 4.

**Figure 4.**
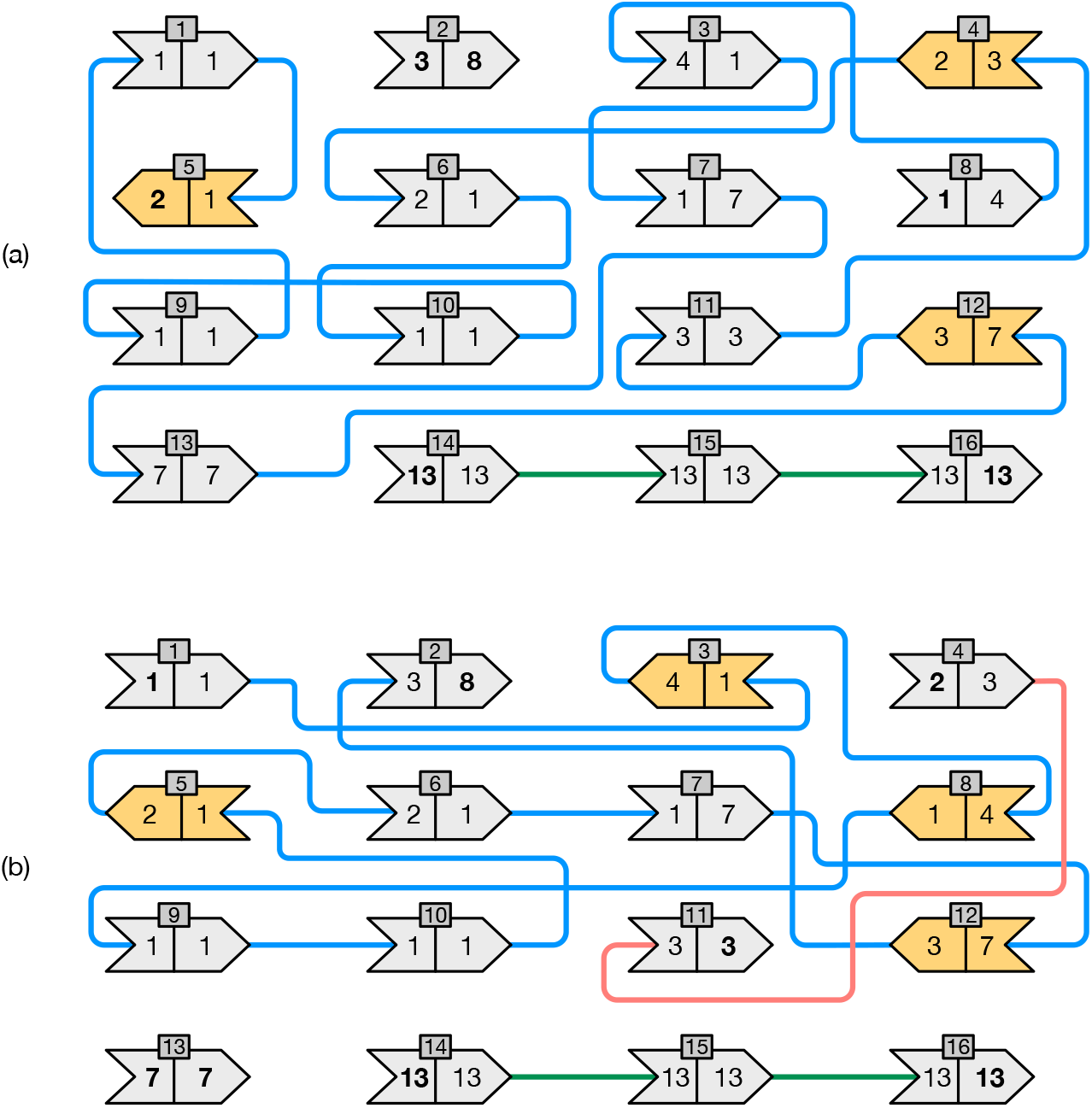
Two different path covers for an example graph with 16 nodes. Nodes linked by edges with the same color belong to the same cover; yellow nodes are those whose orientation was changed. For the graph in the picture, the lower bound in Corollary 9 yields |*C*| *≥* 3. The covers in (a) contains 3 paths and is, therefore, optimal. The cover in (b), instead, contains 4 paths.

Lastly, we summarize the main result of this section with the following Theorem.

##### Theorem 12.

Let 𝒮 be a collection of *m* weighted strings. Let *eW* be the set of the distinct end-point weights of the strings in 𝒮 and 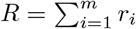, where *r*_*i*_ is the number of runs in the weights of the *i*-th string of 𝒮. Then 𝒮 can be permuted to form at least

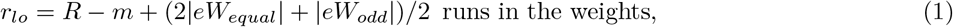

where 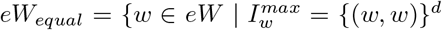, for some *d >* 0} and 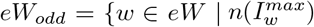 is odd} are defined using the incidence sets for the nodes of *ewG*(𝒮).

**Proof**. There are exactly *R* − *m* runs in 𝒮 that do not depend on the order of the strings in 𝒮. Then 𝒮 can be permuted as to have *R* − *m* + |*C*| runs where *C* is a disjoint-node path cover for *ewG*(𝒮). The theorem follows for Corollary 9 on *C*. ◄

#### 4.2 Computing a Cover

We showed, via a linear-time reduction (Algorithm 2), that the problem of finding a permutation 𝒮 for with the goal of reducing the number of runs in the weights is equivalent to that of computing a path-cover *C* for the graph *ewG*(𝒮). In Section 4.1 we also gave a lower bound to the number of runs that our strategy can achieve. The lower bound depends on the number of paths in *C* (Corollary 9). In this section we therefore present an algorithm to actually compute *C* in *expected linear-time* in the number of nodes of *ewG*(𝒮). Recall that *ewG*(𝒮) has *m* = | 𝒮| nodes, so the complexity is Θ(*m*).

The algorithm is given in Algorithm 3. It manipulates the incidence sets for the end-point weights and a set of unvisited nodes, respectively indicated with *incidence* and *unvisited* in the pseudo-code, and initialized in the lines 4-7. The main loop in the lines 8-22 takes an arbitrary unvisited node and starts a new path from that node. The inner loop in the lines 11-20 greedily tries to extend the current path as much as possible: at every step of the loop, (1) a node is appended to the path, (2) it is erased from the set of unvisited nodes and from the incidence sets where it belongs to, then (3) the next node to append is selected from one of the two incidence sets for the path’s end-point weights. When no extension is possible for both ends, the current path is printed in the lines 21-22. In practice, the output can be written to a file, from which the signed permutation *π* can be derived with Algorithm 2.

##### Algorithm 3

The `cover` algorithm takes an input end-point weight graph *ewG*(𝒮) and prints a set of paths covering the nodes of *ewG*(𝒮).

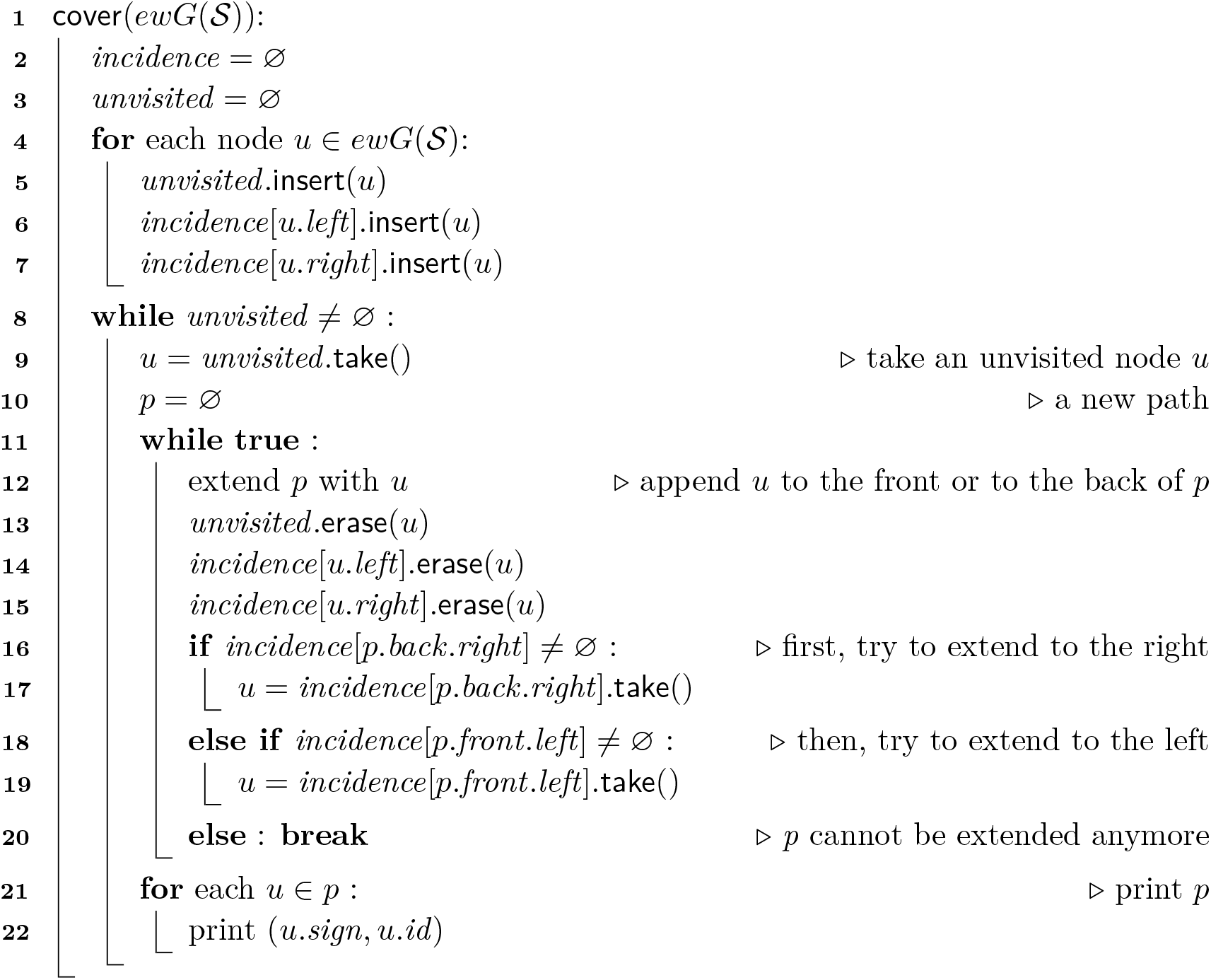

If we use hashing to implement the sets *incidence* and *unvisited*, then the operations `insert`/`erase`/`take` are all supported in *O*(1) expected time. Also appending a node to one of the two path’s end-points can be done in constant (amortized) time using a double-ended queue to represent a path. Figure 5 illustrates how the path is extended with a node (line 12). Therefore, the algorithm runs in expected Θ(*m*) time and consumes Θ(*m*) space because: (1) at most 2*m* nodes (and at least *m*) are inserted in *incidence* and exactly *m* in *unvisited* during the initialization lines 4-7; (2) during the main loop in the lines 8-22, each node is visited, appended to a path, and printed exactly once^2^.

**Figure 5.**
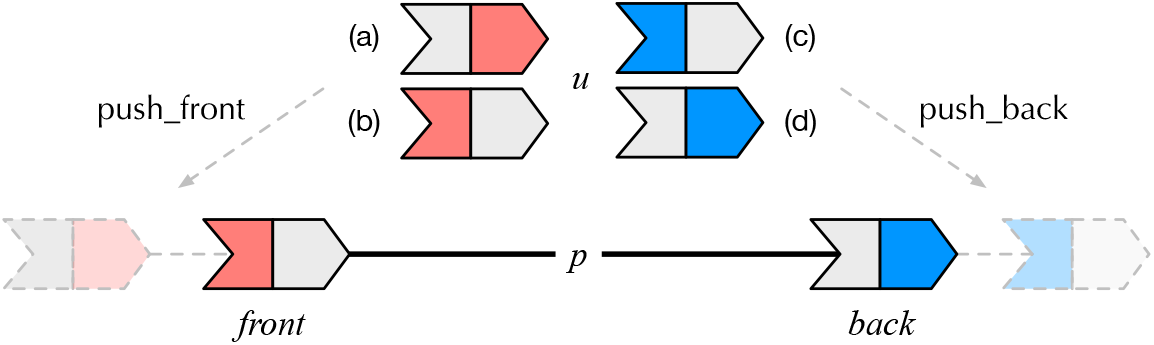
A graphical visualization of line 12 in Algorithm 3 which extends the current path *p* with a node *u*. When *p* is not empty, four different cases can arise, as illustrated in (a), (b), (c), and (d). In cases (b) and (d), the sign of *u* is changed to match one of the two path’s end-points.

## 5 Experiments

In this section we evaluate the proposed weight compression scheme for SSHash and compare it to several competitive baselines. We first describe our experimental setup.

Experiments were run using a server machine equipped with an Intel i9-9940X processor (clocked at 3.30 GHz) and 128 GB of RAM. All the tested software was compiled with gcc 11.2.0 under Ubuntu 19.10 (Linux kernel 5.3.0, 64 bits), using the flags `-03` and `-march=native`. Our implementation of SSHash is written in C++17 and available at https://github.com/jermp/sshash.

All timings were collected using a single core of the processor. The dictionaries are loaded in internal memory before executing queries. For all the experiments, we fix *k* to 31.

### Datasets

We use the following genomic collections: `E-Coli` and `C-Elegans` are, respectively, the full genomes of *E. Coli* (Sakai strain) and *C. Elegans* that were also used in the experimentation by Shibuya et al. [35]; `S-Enterica-100` is a pan-genome of 100 genomes of *S. Enterica*, collected by Rossi et al. [32]; `Human-Chr-13` is the 13-th human chromosome from the genome assembly GRCh38. Table 1 reports some basic statistics for the collections. The weights were collected using the tool BCALM (v2) [7]. In general, note the very low empirical entropy of the weights, *H*_0_(*W*). This is expected since most *k*-mers actually appear once for large-enough values of *k*. Instead, the weights on the pan-genome `S-Enterica-100` have much higher entropy due to the fact that many *k*-mers have weight equal to the number of genomes in the collection (in this specific case, equal to 100). This is useful to test the effectiveness of our encoding on both low- and high-entropy inputs.

**Table 1.** Some basic statistics for the datasets used in the experiments, for *k* = 31, such as: number of distinct *k*-mers (*n*), number of distinct weights (|𝒟|), largest weight (*max*), expected weight value (*E*), and empirical entropy of the weights (*H*_0_(*W*)).

| Dataset | $n$ | $\vert \mathcal{D}\vert $ | $\lceil \log_2 \vert \mathcal{D}\vert \rceil$ | $max$ | $\lceil \log_2 max \rceil$ | $E$ | $H_0(W)$ |
| --- | --- | --- | --- | --- | --- | --- | --- |
| E-Coli | 5,235,781 | 22 | 5 | 27 | 5 | 1.05 | 0.206 |
| S-Enterica-100 | 13,074,614 | 587 | 10 | 3,483 | 12 | 37.47 | 4.420 |
| Human-Chr-13 | 90,911,778 | 806 | 10 | 6,354 | 13 | 1.08 | 0.160 |
| C-Elegans | 94,006,897 | 398 | 9 | 3,478 | 12 | 1.07 | 0.223 |

### Weight Compression in SSHash

We now consider the space effectiveness of the encoding scheme described in Section 3. Table 2 reports the space as average bits per *k*-mer: we see that, in all cases, the space is well below the empirical entropy lower bound *H*_0_(*W*) – usually below by several times. The optimization strategy described in Section 4 brings further advantage. (The space shown is comprehensive of the |𝒟|⌈log_2_ *max*⌉ bits used to represent the distinct weights in the collection. Note that this space takes a negligible fraction of the total space since |𝒟| is very small as reported in Table 1.)

**Table 2.** Space for the weights in SSHash reported in bits/*k*-mer, *before* and *after* the run-reduction optimization from Section 4. For reference, we also report how many times the achieved space is better than the empirical entropy of the weights *H*_0_(*W*).

| Dataset | $H_0(W)$ | *before* | | *after* | |
| --- | --- | --- | --- | --- | --- |
| E-Coli | 0.206 | 0.017 | (12.11×) | 0.014 | (15.10×) |
| S-Enterica-100 | 4.420 | 0.592 | (7.47×) | 0.401 | (11.02×) |
| Human-Chr-13 | 0.160 | 0.136 | (1.18×) | 0.107 | (1.50×) |
| C-Elegans | 0.223 | 0.069 | (3.23×) | 0.055 | (4.05×) |

Table 3, instead, shows the performance of the path cover Algorithm 3. As already mentioned in Section 3, the set of strings indexed by SSHash is obtained by building a spectrum-preserving string set (SPSS) from the raw genome, using the algorithm UST [29] over the output of BCALM [7]. (At our code repository https://github.com/jermp/sshash we provide further details on how to take these preliminary steps before indexing with SSHash.) The number of strings in each collection, *m*, determines the run-time of Algorithm 3 whose complexity is Θ(*m*). The linear-time complexity is evident from the reported timings and makes the algorithm very fast, taking on average a fraction of a microsecond per node.

**Table 3.** The number of input strings (*m*) for SSHash as computed by UST [29], the lower bound on the number of runs (*r*_*lo*_) computed using Equation (1) in Theorem 12, and number of actual runs (*r*) after the optimization. We report the increase, in percentage, of *r* compared to *r*_*lo*_. The last two columns show the run-time of the path cover Algorithm 3, in total milliseconds (ms) and average nanoseconds per node (ns/node).

| Dataset | $m$ | $r_{lo}$ | $r$ | | Alg. 3 (ms) | Alg. 3 (ns/node) |
| --- | --- | --- | --- | --- | --- | --- |
| E-Coli | 2,102 | 3,723 | 3,723 | (+0.0000%) | 0.6 | 285 |
| S-Enterica-100 | 150,604 | 277,649 | 277,658 | (+0.0032%) | 53.0 | 352 |
| Human-Chr-13 | 266,113 | 462,175 | 462,197 | (+0.0048%) | 94.6 | 355 |
| C-Elegans | 140,452 | 247,661 | 247,669 | (+0.0032%) | 47.1 | 335 |

The other important point to note is that Algorithm 3 is empirically *optimal* regarding the number of runs given that the final number of runs, *r*, is only slightly higher than the lower bound *r*_*lo*_ computed using Equation (1).

(In Appendix B we report additional experimental results.)

### Overall Comparison

In Table 4 we show a comparison between the following weighted dictionaries (see also Section 2; links to the code repositories are included in the References):

**Table 4.**
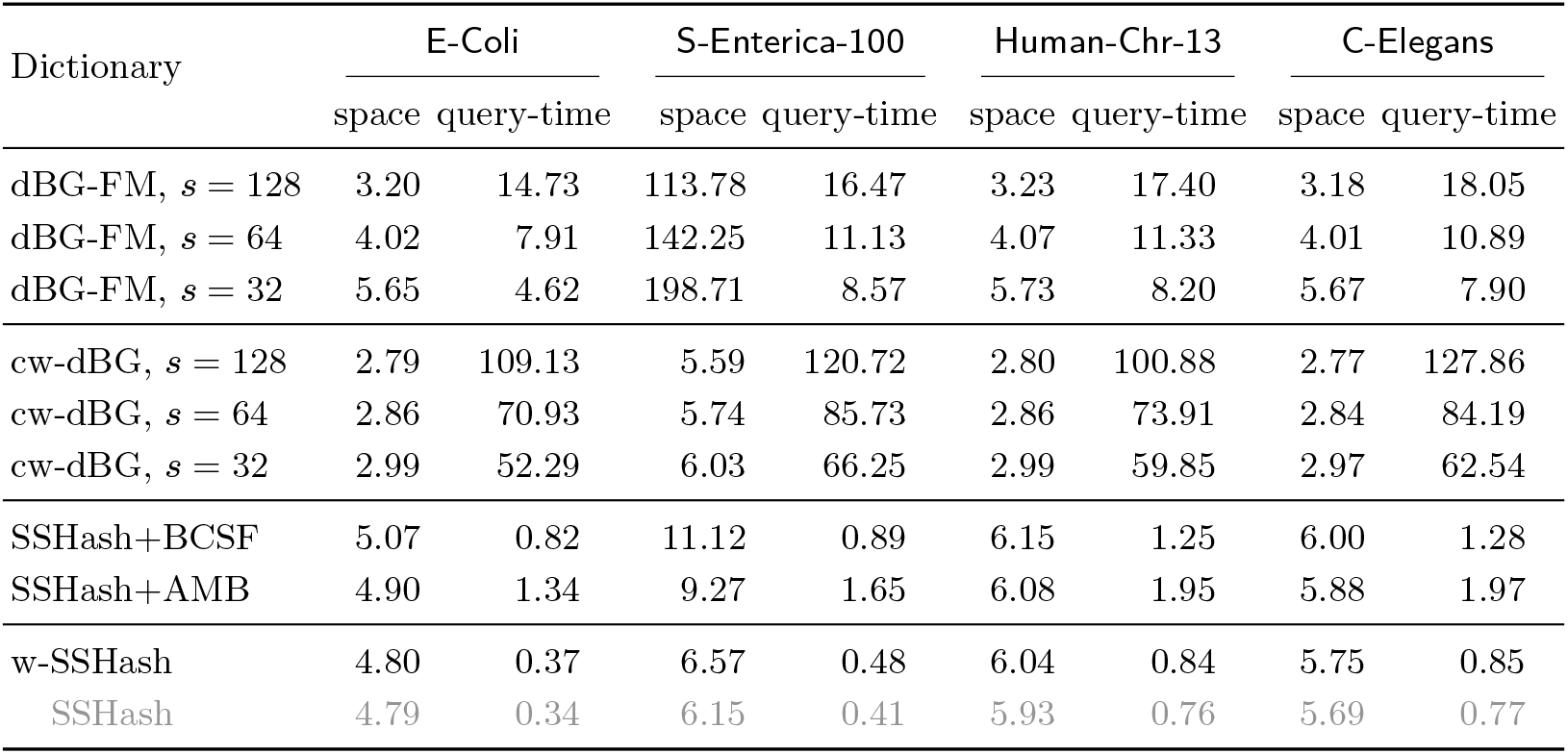
Dictionary space in average bits/*k*-mer and count time in average *μ*sec/*k*-mer. For reference, we report in gray color the space and time of SSHash *without* the weight information.

▬ The dBG-FM index [6] based on the popular FM-index [11]. In particular, this representation implements a weighted *k*-mer dictionary via the *count* query which returns the number of occurrences of a given *k*-mer in the input. The *count* query, in turn, is implemented using rank queries over the BWT. The dBG-FM implementation has a main trade-off parameter, *s*, to control the practical performance of rank queries. We test the values *s* = 32, 64, 128.
▬ The recent cw-dBG [12] dictionary based on the data structure called BOSS [4]. Similarly to an FM-index, also cw-dBG has a trade-off parameter that we vary as *s* = 32, 64, 128. (The authors used *s* = 64 in their own experiments.)
▬ The *non*-weighted SSHash itself coupled with the fast compressed static function (CSF) tailored for low-entropy distributions, proposed by Shibuya et al. [35]. As reviewed in Section 2, a CSF does not represent the *k*-mers but just realizes a map from *k*-mers to their weights. Such map is collision-free only over the set of *k*-mers that was used to actually build the function. Therefore, we use SSHash as an efficient dictionary for the *k*-mers and the CSF to represent the weights. The authors proposed two different versions of their approach, BCSF and AMB, with different space/time trade-offs.
▬ The weighted SSHash dictionary proposed in this work, which we refer to as w-SSHash in the following, *after* the run-reduction optimization (Table 2 and 3). We use the *regular* index variant of SSHash. The main parameter of the index – the *minimizer* length – is always set to ⌈log_4_ *N*⌉ + 1 where *N* is the number of nucleotides in the SPSSs of the datasets, following the recommendation given in the previous paper [22]. Therefore, we use the following minimizer lengths: 13, 14, 15, and 15, for respectively, `E-Coli`, `S-Enterica-100`, `Human-Chr-13`, and `C-Elegans`. Also the AMB algorithm by Shibuya et al. [35] is based on minimizers and we use the same lengths.

We did not compare against deBGR [19] and Squeakr [21] as the authors of cw-dBG showed in their experimentation [12] that both tools take considerably more space than cw-dBG, e.g., one order of magnitude more space. Here, we are interested in a good balance between space effectiveness and query efficiency.

To measure query-time – the time it takes to retrieve the weight *w*(*g*) given the *k*-mer *g* – we sampled 10^6^ *k*-mers uniformly at random from the collections and use them as queries. We report the mean between 5 measurements. Half of the queries were transformed into their reverse complements to make sure we benchmark the dictionaries in the most general case.

The space of w-SSHash is generally competitive with that of the fastest variant of dBG-FM (*s* = 32), but w-SSHash has (more than) one order of magnitude better query time. Note that on `S-Enterica-100` the dBG-FM index is space-inefficient since it redundantly represents many repeated *k*-mers. Using a higher sampling rate reduces the space of dBG-FM at the price of slowing down query-time; however, the most space-efficient variant tested (*s* = 128) is not even 2*×* smaller than w-SSHash.

The cw-dBG index is the smallest tested dictionary. Its space effectiveness is comparable to that of dBG-FM *s* = 128, and indeed generally twice as better as that of w-SSHash. The price to pay for this enhanced compression ratio is a significant penalty at query-time. Indeed, w-SSHash can be two order of magnitude faster than cw-dBG. Consider, for example, the two dictionaries built for `S-Enterica-100`: we have 0.5 vs. 66-120 *μ*s per query.

The two CSFs, BCSF and AMB, make SSHash 2-3*×* slower than w-SSHash and even consistently larger. This comparison motivates the need for a unified data structure to handle efficiently both the *k*-mers *and* the weights, like w-SSHash. While the increase in space due to the CSF is not much for the low-entropy datasets because both BCSF and AMB are very space-efficient in those cases, the gap is more evident on `S-Enterica-100`.

As a last note, observe that there is no significant slowdown in accessing the weights in w-SSHash compared to a simpler membership query (the time reported in shaded color in Table 4), hence proving the RLE-based scheme to be efficient too and not only very effective.

## 6 Conclusions

In this work we extended the recent SSHash [22] dictionary to also store the weights of the *k*-mers in compressed format. In particular, we represented the weights using compressed *runs* of equal symbols. While using run-length encoding to compress highly repetitive sequences is not novel per se and indeed a folklore strategy at the basis of many other data structures, this allows to use a very small extra space (e.g., much less than the empirical entropy of the weights) on top of SSHash with only a slight penalty at retrieval time. The crucial point is that it is possible to use run-length encoding because SSHash *preserves the (relative) order* of the *k*-mers in the indexed sequences. The main practical take-away is, therefore, that SSHash handles weighted *k*-mer sets in an *exact* manner without noticeable extra costs. Our software is publicly available to encourage its use and reproducibility of results.

We also introduced the concept of *end-point weight graph* (*ewG*) and showed its usefulness in reducing the number of runs in the weights. Precisely, we showed that minimizing the number of runs in a collection of sequences corresponds to the problem of computing a minimum-cardinality path cover for the *ewG* of the sequences. We presented a greedy algorithm that computes a cover in expected linear-time (in the number of nodes of the graph) and showed that it is empirically almost optimal according to a lower bound on the number of runs. As a result of this optimization, the space spent to represent the weights is unlikely to be improved using run-length encoding.

Although several approaches in the literature [17, 21, 35, 36] also consider *approximate* weights, we did not pursue this direction here as the weights are already encoded space-efficiently in SSHash and in an *exact* way, so there may be no need for approximation.

The distribution of weights in large collections is usually expected to be very skew, i.e., most *k*-mers actually appear once and few of them repeat many times [35, 36]. A common strategy to save space is then to avoid the representation of the most frequent weight(s). Note that, since we represent runs of weights and not the individual weights, we are already optimizing (potentially very large) sub-sets of weights equal to the most frequent one. That is, run-length encoding is also a good match for such skew distributions.

## Supplementary Material

Source code: https://github.com/jermp/sshash.

## A Omitted Proofs from Section 4.1

### Lemma 3.

**Proof**. We proceed by induction on *d*. Base case: if *d* = 1 (odd case), then there is only the singleton path (*w, x*); if *d* = 2 (even case), then we can either form the path (*w, x*) → (*x, w*) of end-points (*w, w*) or the path (*x, w*) → (*w, x*) of end-points (*x, x*). So the base case is verified. Now we assume the Lemma holds true for a generic *d >* 2 and we want to prove it for *d* + 1. If *d* is even, then *d* + 1 is odd and we can either have a path (*w, w*) → (*w, x*) or a path (*w, x*) → (*x, x*). In both cases the end-points are (*w, x*). Symmetrically: if *d* is odd, then *d* + 1 is even and we can either have a path (*w, x*) → (*x, w*) with end-points (*w, w*) or a path (*x, w*) → (*w, x*) with end-points (*x, x*).

### Lemma 6.

**Proof**. Let us first consider the special case where all the other weights in *I*_*w*_ are distinct, so there are no equal nodes in *I*_*w*_ except for, possibly, nodes of the form (*w, w*). In this case, we say that *I*_*w*_ is *canonical*. If there are some nodes (*w, w*), they can be trivially collapsed to a single path of end-points (*w, w*) by Lemma 3. So, without loss of generality, either we have *one* node (*w, w*) in *I*_*w*_ and *n*(*I*_*w*_) = |*I*_*w*_| + 1, or we do not and *n*(*I*_*w*_) = |*I*_*w*_|. In this special case, since *n*(*I*_*w*_) is odd, it is always possible to create 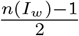 paths, each having 2 nodes. These paths will *not* contain *w* as end-point because all the end-points where *w* appears are used to link the nodes. Therefore, there will be exactly one unpaired node where *w* appears.

Now, observe that we can relax the restriction on the other weights to be all distinct. For every weight *x* ≠ *w* that appears for *d*_*x*_ *>* 1 times in *I*_*w*_, there are *d*_*x*_ equal nodes (*w, x*). Let *D*_*x*_ ⊆ *I*_*w*_ be the set of such nodes (hence, *d*_*x*_ = |*D*_*x*_|). By applying Lemma 3 to the nodes of each set *D*_*x*_:

▬ If *d*_*x*_ is even, then the nodes in *D*_*x*_ can be collapsed into a path of end-points (*w, w*) or (*x, x*). If the node (*w, w*) is created, we obtain a new incidence set 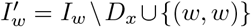. Instead, if the node (*x, x*) is created, then 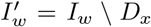 since (*x, x*) cannot be in an incidence set for *w*. In both cases 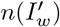 will still be odd since we subtract an even number from *n*(*I*_*w*_).
▬ If *d*_*x*_ is odd, the nodes are collapsed into the path of end-points (*w, x*) and the new incidence set is 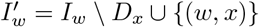. Again, 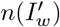 will still be odd since we subtract an odd number from *n*(*I*_*w*_) but sum one.

After each set *D*_*x*_ is processed in this way, we are left with an incidence set for *w* that is canonical. ◄

### Lemma 8.

|*W*_*odd*_| is even.

**Proof**. Observe that 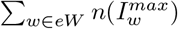 is even and equal to 2*m* because we count the occurrences of the weights appearing as end-points of the sequences and each sequence has two end-points. Since *eW* = *eW*_*odd*_ ∪ *eW*_*even*_ ∪ *eW*_*equal*_, the above sum can be re-written as

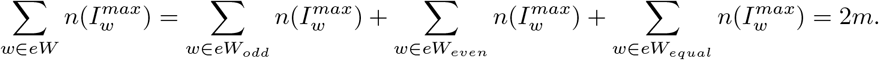

It follows that also

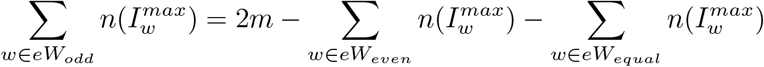

must be even since it is obtained by difference of even quantities. Since each term in the sum 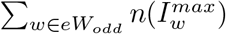 is odd by definition, the whole sum is even if and only if |*eW*_*odd*_| is even, as the sum of an odd number of odd numbers is odd. ◄

## B Additional Experimental Results

In Table 5 and Table 6 we report the performance of Alg. 3 and of w-SSHash on four additional, larger, collections that we also used in our previous work [22], namely the full genomes of *G. Morhua* (`Cod`), *F. Tinnunculus* (`Kestrel`), and *H. Sapiens* (`Human`), and a collection of more than 8000 bacterial genomes (`Bacterial`) [1]. Precisely, the results in Table 6 are for regular w-SSHash dictionaries with minimizer lengths equal to 17, 17, 20, and 20, for respectively, `Cod`, `Kestrel`, `Human`, and `Bacterial`.

**Table 5.** The performance of Alg. 3 on the datasets Cod, Kestrel, Human, and Bacterial, for which we report the number of distinct *k*-mers (*n*) and the number of strings (*m*) after running UST [29] on the collections. The performance of the algorithm is expressed as: the number of actual runs (*r*) after the run-reduction optimization in comparison with the lower bound on the number of runs (*r*_*lo*_) computed using Equation (1), and running time (in total sec and average ns/node).

| Dataset | $n$ | $m$ | $r_{lo}$ | $r$ | | Alg. 3<br>(sec) | Alg. 3<br>(ns/node) |
| --- | --- | --- | --- | --- | --- | --- | --- |
| Cod | 502,465,200 | 2,406,681 | 4,183,202 | 4,183,230 | (+0.00067%) | 1.2 | 500 |
| Kestrel | 1,150,399,205 | 682,344 | 1,140,743 | 1,140,747 | (+0.00035%) | 0.3 | 440 |
| Human | 2,505,445,761 | 13,014,641 | 22,680,047 | 22,680,099 | (+0.00023%) | 7.5 | 580 |
| Bacterial | 5,350,807,438 | 26,448,286 | 56,662,230 | 56,662,304 | (+0.00013%) | 17.2 | 650 |

**Table 6.**
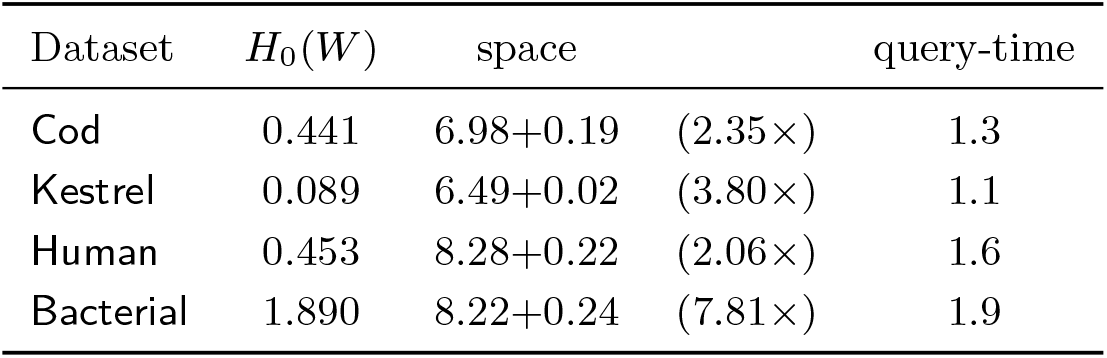
The performance of w-SSHash on the permuted string collections Cod, Kestrel, Human, and Bacterial. We report the empirical entropy of the weights (*H*_0_(*W*)), the dictionary space in average bits/*k*-mer, and query-time in average *μ*sec/*k*-mer. The space is indicated as *x* + *y*, where *x* is the space of SSHash (without the weights) and *y* is the space for the encoding of the weights.

## Footnotes

1 Elias-Fano represents a monotone integer sequence *S*[1..*n*] with *S*[*n*] ≤ *U* in at most *n*⌈log_2_(*U/n*)⌉ + 2*n* bits. With *o*(*n*) extra bits it is possible to decode any *S*[*i*] in constant time and support predecessor queries in *O*(log(*U/n*)) time. For a complete description of the method, we point the reader to the survey by Pibiri and Venturini [28, Section 3.4]. We also remark that Elias-Fano has been recently used in many compressed, practical, data structures (see, e.g., [25, 26, 27]).

2 An important implementation remark: In practice, we implemented the *incidence* sets using a single open-addressing hash-table with a sorted array of 2*m* nodes and an extra array of offsets into the vector to distinguish between the different incidence sets. This makes the algorithm much faster and lighter than a straightforward implementation using a linear-chaining hash-table (e.g., the C++’s `std::unordered_set`), with chains usually implemented as linked lists.

